# Evidence of decreased gap junction coupling between astrocytes and oligodendrocytes in the anterior cingulate cortex of depressed suicides

**DOI:** 10.1101/578807

**Authors:** Arnaud Tanti, Pierre-Eric Lutz, John Kim, Liam O’Leary, Gustavo Turecki, Naguib Mechawar

## Abstract

Glial dysfunction is a major feature in the pathophysiology of mood disorders. While altered astrocyte (AS) and oligodendrocyte-lineage (OL) cell functions have been associated with depression, the crosstalk between these two major glial cell types has never been assessed in that context. AS are potent regulators of OL cells and myelination, in part through gap junction-mediated intercellular communication made possible by the heterotypic coupling of AS-specific (Cx30 and Cx43) and OL-specific (Cx32 and Cx47) connexins, allowing cytosolic transport and metabolic support to OL cells. Because changes in the expression of AS-specific connexins have been previously reported in the brain of depressed individuals, this study aimed at addressing the integrity of AS-OL coupling in the anterior cingulate cortex (ACC) of depressed suicides. Using immunofluorescence and confocal imaging, we characterized the distribution of the AS-specific Cx30 in the ACC, and mapped its expression onto oligodendrocyte somas and myelinated axons as well as brain vasculature in post-mortem brain samples from depressed suicides (N=48) and matched controls (N=23). The differential gene expression of key components of the gap junction nexus was also screened through RNA-sequencing dataset previously generated by our group, and validated by quantitative real-time PCR. Our results indicate that Cx30 expression mapping to OL cells is selectively decreased in depressed suicides, an effect that was associated with decreased expression of OL-specific connexins Cx32 and Cx47, as well as the downregulation of major connexin-interacting proteins essential for the scaffolding, trafficking and function of gap junction channels. These results provide a first evidence of impaired gap junction mediated communication between astrocytes and oligodendrocytes in the ACC of individuals with mood disorders. These changes in glial coupling are likely to have significant impact on brain function, and may contribute to the altered OL function previously reported in this brain region.

## INTRODUCTION

Glial dysfunction has emerged as an important pathophysiological hallmark of depression and suicide. In particular, post-mortem studies have consistently shown changes in astrocyte numbers and morphology, as well as in the expression of several astrocytic markers in fronto-limbic regions of depressed individuals^1–10^, while pharmacological or genetic disruption of astrocyte function in rodents has been associated with depressive-like behaviour^11,12^. Recently, oligodendrocyte-lineage (OL) cells have also been associated with depression and suicide. These cells, which allow for a formidable form of brain plasticity by responding to environmental cues and experiences in an activity-dependent manner^13–15^, were found to display signs of functional impairments in both post-mortem brain samples from depressed individuals^16–24^ and stress-induced animal models of depression^25–27^. These findings could represent a neurobiological substrate to the altered connectivity and white matter integrity that have been reported in imaging studies^28–35^, highlighting that myelin plasticity in discrete cortico-limbic areas could mediate some of the behavioral changes characterizing depression or modulate individual vulnerability to this mood disorder.

While astrocytes and oligodendrocytes have been independently implicated in the pathophysiology of depression, the interplay between these cell types has never been examined in this context. Astrocytes are known potent regulators of OL cell proliferation, survival and maturation as well as myelination. This multi-faceted influence of astrocytes is exerted either indirectly through their implication in neurotransmission, or directly through the release of various molecules affecting the physiology of oligodendrocytes and myelin maintenance^36–39^.

Part of the astrocytic control over OL cells and myelination is permitted through cytosolic exchanges allowed by gap junction coupling between astrocytes and oligodendrocytes^36,38,40,41^. Gap junction coupling, which can also be homotypic, thus gives rise to extended networks of interconnected glial cells in the brain. The docking of two connexin hexamers (hemichannel, or connexon) from two adjacent cells forms a channel allowing the exchange of a wide range of small molecules such as ions, metabolites, second messengers and neurotransmitters, referred to as gliotransmitters^42^. Astrocytes mainly express Cx30 and Cx43, while oligodendrocytes mainly express Connexin 32 (Cx32) and Cx47. Functional channels are formed between astrocytes and oligodendrocytes (A:O) through the heterotypic coupling of Cx30-Cx32 and Cx43-Cx47^41,43–47^.

Glial gap junction coupling has been shown to be of critical importance for oligodendrocyte function, including normal myelination. In double knockout mouse models, loss of oligodendrocyte-expressed Cx32 and Cx47 induces important abnormalities in the central nervous system^48^, including thinner or absent myelin sheaths, oligodendrocyte cell death and axonal loss. Interestingly, the double deletion of both astrocyte-expressed Cx30 and Cx43 also leads, albeit to a lesser extent, to myelination impairments^49^. Moreover, mice with concomitant deletions of an oligodendrocyte connexin and an astrocyte connexin, for example Cx43/Cx32 or Cx30/Cx47 double knock-out mice, show impaired astrocyte-oligodendrocyte coupling, severe myelin pathology such as myelin loss, and decreased numbers of OL cells^50,51^. Overall, these studies demonstrate that astrocyte-oligodendrocyte gap junction coupling is particularly important for oligodendrocyte function.

Our group and others recently reported that the expression of both astrocytic Cx30 and Cx43 is significantly decreased in multiple areas of the depressed suicide brain, including the prefrontal cortex, orbitofrontal cortex, mediodorsal thalamus, and caudate nucleus^7,52^. These findings have been replicated in chronic stress mouse models of depression^53,54^, with decreased diffusion of gap junction permeable dyes concomitant to changes in Cx43 expression^54^. The underlying mechanisms and functional consequences of such coupling changes are unclear, but given that astroglial networks are involved, notably, in regulating neuronal activity and plasticity, metabolic and osmotic support, blood-brain barrier integrity, and myelination^38,55–57^, alterations in glial coupling are likely to have significant consequences on brain function. Importantly, it remains to be determined whether decreased expression of astrocytic connexins may reflect changes in gap junction coupling of astrocytes among themselves, and/or with other cell types.

We examined this question in the present study by focusing on astrocyte-oligodendrocyte coupling in post-mortem anterior cingulate cortex (ACC) samples from depressed suicides and matched sudden-death controls. This cortical region was selected because we previously found it to display convergent molecular and cellular abnormalities of myelination and oligodendrocyte function in depressed suicides^18^, particularly in individuals with a history of child abuse (CA). Based on these previous results and evidence suggesting that such individuals may represent a distinct clinical and neurobiological subtype of depression^58^, we also examined samples from a group of depressed suicides with a history of CA. Our experimental strategy was to combine double-labeling immunofluorescence and confocal microscopy to map the expression of the astrocyte connexin Cx30 onto oligodendrocyte somas and myelinated fibers. We also mapped Cx30 expression onto cerebral vasculature as a way to address the cell-type specificity of the potential changes observed in Cx30 mapping. We also screened for the expression of genes known to be important for the scaffolding and function of gap junction channels (GJCs), as well as genes coding for the main astrocyte- and oligodendrocyte-specific connexins. These experiments provided evidence of a significant decrease of Cx30-immunoreactivity (-IR) specifically on oligodendrocyte-lineage cells and a significant downregulation of oligodendrocyte-specific connexins, as well as key associated molecules in the ACC of depressed suicides, altogether suggesting impaired AS-OL gap junction communication in the brain of depressed suicides.

## MATERIAL AND METHODS

### Human post-mortem samples

This project was approved by the Douglas Hospital Research Ethics Board. Brain tissues were obtained from the Suicide section of the Douglas-Bell Canada Brain Bank (Douglas Hospital Research Centre, Montreal, Canada) in collaboration with the Quebec Coroner’s Office and with informed consent from next of kin. Samples were obtained from individuals having died by suicide in the context of a depressive episode (Depressed Suicides – DS; N=48) and from healthy controls (Controls - CTRL; N= 23) having died suddenly without psychiatric or neurological illness. Groups were matched for age, post-mortem interval (PMI), tissue pH and RNA Integrity Number (RIN) (**Table 1**). Cases and controls were defined with the support of medical charts and Coroner records, and based on previously described and validated psychological autopsies^59^. In brief, the Structured Clinical Interview for DSM-IV Psychiatric Disorders (SCID-I) was conducted by a trained interviewer with one or more informants best acquainted with the deceased and reviewed by a panel of clinicians to reach consensual diagnosis. Cases with a history of CA all scored highest (1 or 2) in the rating scale of sexual and physical abuse from an adapted version of the Childhood Experience of Care and Abuse (CECA)^60,61^. Histological experiments for Cx30 quantification and colocalization were performed on a subset of matched controls (N=8-11) and depressed suicides (N= 12-17).

**Table 1.** Subjects characteristics

|  | **CTRL** | **DS-nCA** | **DS-CA** |
| --- | --- | --- | --- |
| **N** | 23 | 23 | 25 |
| **Axis 1 diagnosis** | none | MDD (19); DD-NOS (4) | MDD (21); DD-NOS (4) |
| **History of CA** | none | none | 25 |
| **Age (years)** | 46.2 ± 22.2 | 49.3 ± 10.3 | 42.2 ± 14.8 |
| **Sex (M/F)** | 18/5 | 20/3 | 18/7 |
| **PMI (hours)** | 35.4 ± 21.5 | 51.7 ± 29.4 | 42 ± 22.5 |
| **pH** | 6.5 ± 0.3 | 6.6 ± 0.3 | 6.6 ± 0.3 |
| **RIN** | 6.9 ± 0.6 | 6.5 ± 0.3 | 6.8 ± 0.8 |
| **Substance dependence** | 0 | 9 | 8 |
| **AD medication** | 0 | 9 | 8 |

Data represent mean ± SD. AD: antidepressant; DD-NOS: Depressive Disorder Not Otherwise Specified; MDD: Major Depressive Disorder; PMI: post-mortem interval; RIN: RNA Integrity Number.

### Tissue dissections

As previously described^18^, ACC samples were dissected by expert brain bank staff from frozen coronal sections with the guidance of a human brain atlas^62^. Histology blocks (∼1cm^3^) were collected in sections equivalent to plate 6 of this atlas (−30mm from the center of the anterior commissure) immediately rostral to the genu of the corpus callosum, using the cingulate and the callosal sulci as the rostral and caudal landmark limits, respectively. Frozen blocks were fixed overnight in 10% formalin, cryoprotected in a 20% sucrose/PBS solution, and cut serially in 50µm-thick sections with a cryostat. Sections were kept in cryoprotectant (30% glycerol, 30% ethylene glycol, PBS) until processed for immunofluorescence.

### Immunohistochemistry

Free-floating sections were rinsed in PBS and then incubated overnight and at room temperature in primary antibodies diluted in a solution of PBS-Tx-100/2% normal donkey serum. Following washes in PBS, sections were incubated for 2 hours at room temperature with Alexa-488, Cy3 or Cy5-conjugated secondary antibodies produced in donkey (Jackson ImmunoResearch) diluted 1:500 in PBS-Tx-100/2% normal donkey serum. Sections were then mounted on slides, coverslipped (Vectashield mounting medium with DAPI, Vector Laboratories) and kept at 4°C until imaging. The primary antibodies used in this study along with their respective dilutions are listed in **Table 2**.

**Table 2.** Primary antibodies used in this study

| **Primary antibody** | **Species** | **Dilution** | **Manufacturer** | **Reference number** |
| --- | --- | --- | --- | --- |
| Connexin 30 | Mouse | 500 | Santa Cruz Biotechnology | sc-514847 |
| Connexin 43 | Rabbit | 500 | Cell Signaling | 3512 |
| Myelin Proteolipid Protein (PLP) | Rabbit | 1000 | Abcam | ab105784 |
| Nogo-A | Rabbit | 100 | Millipore | AB5888 |
| Laminin | Rabbit | 1000 | Sigma-Aldrich | L9393 |
| Vimentin | Rabbit | 1000 | Abcam | ab92547 |
| GFAP | Chicken | 1000 | Abcam | ab4674 |

### Confocal imaging and analysis

Image acquisitions was performed on an Olympus laser scanning confocal microscope (FV1200) equipped with a motorized stage. For the analysis of Cx30 density and colocalization experiments, stack images were taken with a x60 objective (NA = 1.42) and 3.5 digital zoom, with XY pixel width of 0.06µm and z spacing of 0.8µm. This spacing was chosen to prevent any dot from being counted twice in two successive z slices. Parameters were kept consistent across all acquisitions. For Cx30 total density measurements, 4 random stacks with an average of 15 z-sections per stack were randomly acquired within layer 5/6 of the ACC. For each z-section, the number of Cx30 puncta was quantified by the *find maxima* function of ImageJ with a noise tolerance set at 350, and divided by the image area to calculate a density per mm^2^. We used this approach as it gave the most accurate estimate of puncta number compared to manual counting on a small set of captured images. For each stack the z-sections densities were then averaged, and stack averages computed per subject to yield final Cx30 densities. For Cx30 mapping onto Nogo-A positive cells, an average of 20 cells were manually segmented within 3 images stacks acquired within layer 5/6 of the ACC and using previously described acquisition parameters. For each z-section, cell selection was enlarged by 0.5um, and number of Cx30 puncta quantified as described above, using the *find maxima* function with a noise tolerance of 350. Total Cx30 puncta numbers per cell were averaged to yield the average number of Cx30 mapping to Nogo-A+ cells per subject. Cx30 mapping to myelinated fibers and blood vessels was performed similarly in 4 random stacks taken within layer 5/6 of the ACC and using the same acquisition parameters. For each z-section of the stacks, PLP or Laminin IR were subjected to auto-thresholding and converted to mask. A median filter with a 3 pixels radius was applied, followed by a round of dilation and erosion. For each z-section the number of Cx30 puncta within PLP or Laminin masks was then quantified using the *find maxima* function (noise tolerance of 350), and the density of Cx30 calculated by dividing the number of Cx30 puncta by PLP or Laminin area respectively. Densities per stack were then averaged to yield the final density of Cx30 mapping to PLP or laminin per subject.

### RNA-sequencing and RT-PCR validation

The RNA sequencing dataset, which was generated with a cohort including all of the subjects from the present study, was thoroughly described in a report we recently published^18^. In brief, total RNA was extracted from frozen ACC samples using the RNeasy Lipid Tissue Mini Kit (Qiagen) and a 2100 Bioanalyzer (Agilent) used to measure RNA concentration and RIN. Libraries preparation was performed at the Genome Quebec Innovation Center with the TrueSeq Stranded Total RNA Sample Preparation and Ribo-Zero Gold kits (Illumina), and quantified using PCR (KAPA Library Quantification). Sequencing was done on an Illumina HiSeq 2000 (100bp paired-end), with ∼62 million reads/library. After adapter trimming, alignment and gene quantification using FASTX-Toolkit, Trimmomatic, TopHat v2.1.0 and HTSeq-count respectively, differential expression between groups was analyzed with DESeq2 using RIN, age and gender as covariates. Differential expression was screened in a panel of genes built based on the literature and the evidence of their role in gap junction signaling and cell-cell junctions (**Figure 4**). Validation of differential gene expression was performed by quantitative RT-PCR for the 8 following genes: GJB6, GJA1, GJB1, GJC2, CAV1, CAV2, OCLN and DBN1 (**Table 3**). Starting from 50mg of frozen tissue, total RNA was extracted using RNeasy Lipid Tissue Mini Kit (Qiagen) with DNase digestion. RNA quantity and RIN were measured with the 2200 TapeStation (Agilent), and only samples with RIN>5.5 were used for analysis. One µg of RNA was used to synthetize cDNA using M-MLV reverse transcriptase (Invitrogen) with oligo-dT and random hexamers. Quantitative PCR reactions were performed using SYBR Green intercalating dye and master mix (Biorad) on a QuantStudio 6 Flex Real-Time PCR system (ThermoFisher Scientific), using four replicates per gene. Primers were designed with Primer-BLAST (http://www.ncbi.nlm.nih.gov/tools/primer-blast/) and validated by dissociation curves and gel migration. Relative expression was calculated on the QuantStudio Real-Time PCR Software version 1.2 (ThermoFisher Scientific) using calibration curves and normalization by the housekeeping gene GAPDH.

**Table 3.**
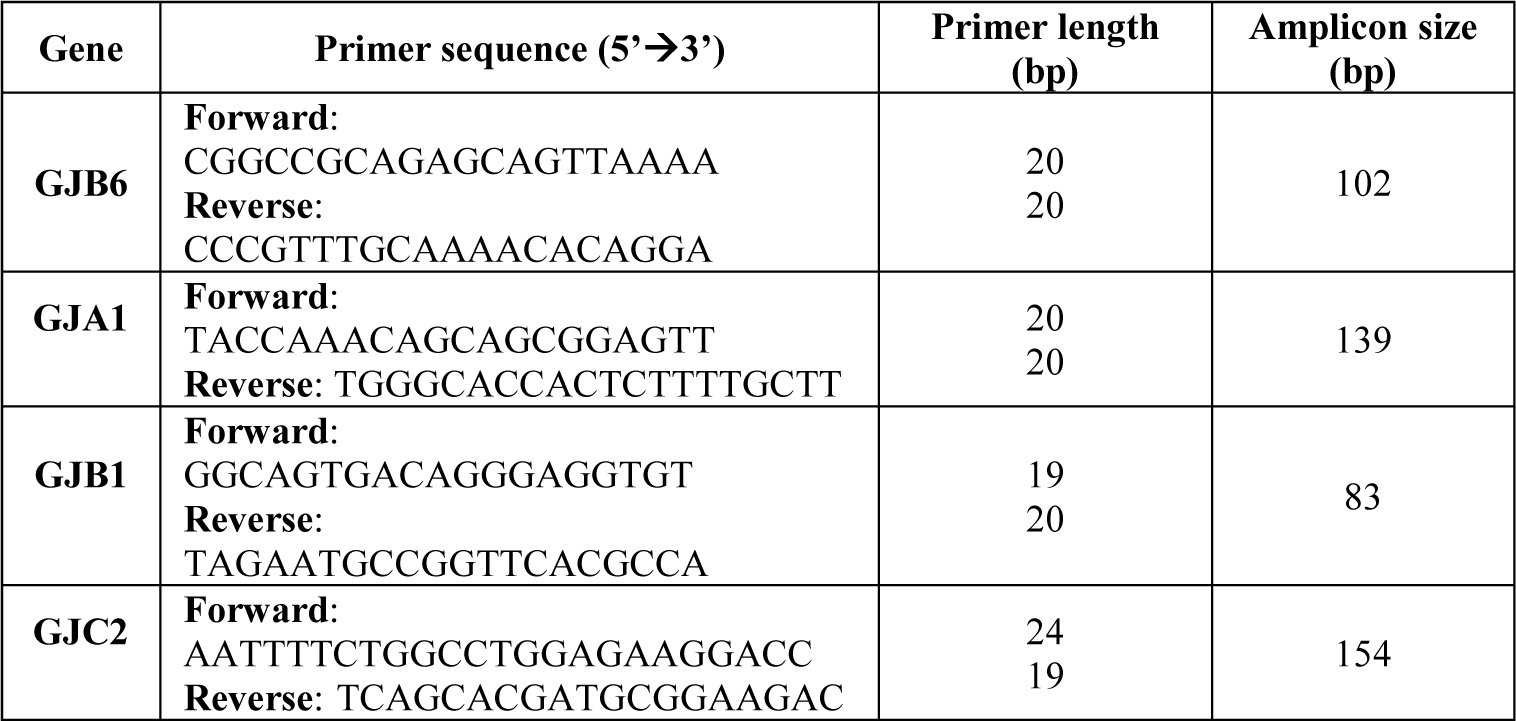
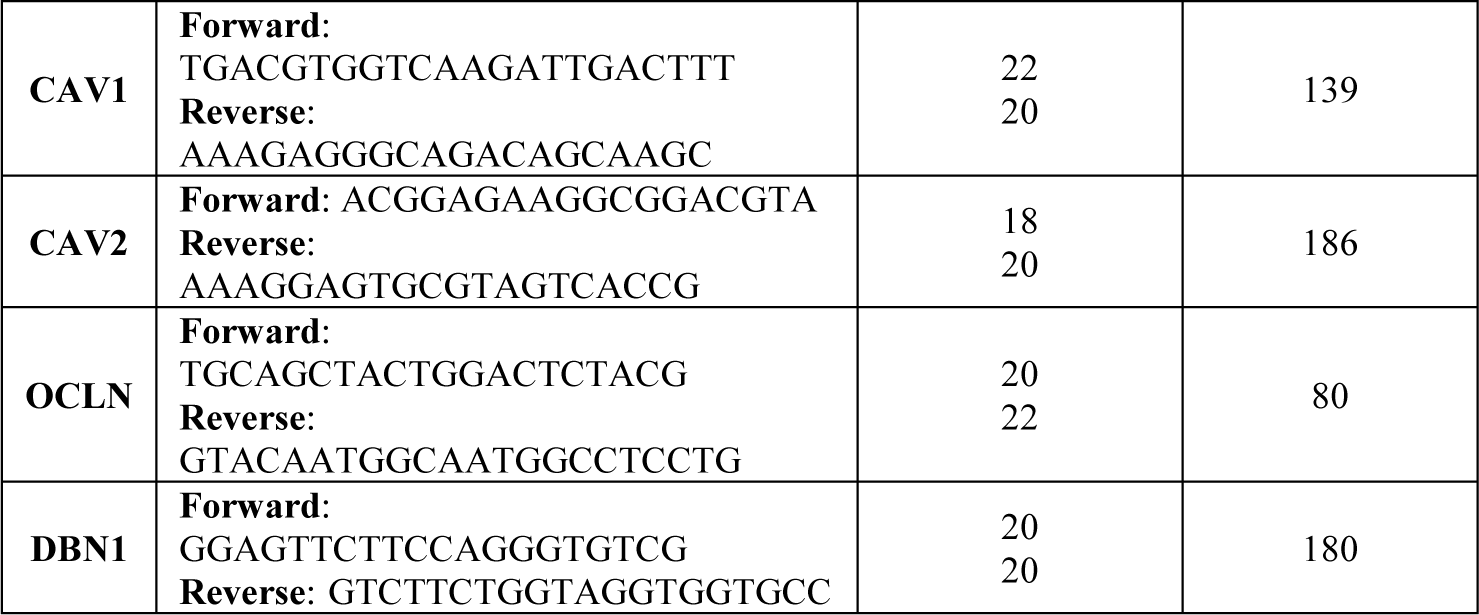
Primers used for quantitative RT-PCR validation of differential expression

### Statistics

All statistical analyses were performed on Statistica version 12 (StatSoft) and Prism version 6 (GraphPad Software). Distribution of the data and homogeneity of variances between groups were assessed with Shapiro-Wilk and Levene’s tests, respectively. For Cx30 density measurements and mapping experiments comparing Controls (CTRL), Depressed Suicides (DS-nCA) and Depressed Suicides with a history of CA (DS-CA), group effects were detected with Kruskal-Wallis non-parametric one-way ANOVAs, followed by Dunn’s multiple comparisons post-hoc test. Effects of a history of substance abuse and medication in Depressed Suicides on Cx30 density were addressed using Kruskal-Wallis one-way non-parametric ANOVA. Linear regressions were used to assess the effects of age and PMI on Cx30 density. Differential expression in the RNA-sequencing dataset was performed using DEseq2, using a linear model controlling for RIN, age and gender. Differential expression in RT-PCR data was assessed using unpaired two-tailed t-tests comparing cases to controls. Nominal p-values are reported with a significance threshold set at 0.05.

## RESULTS

### General distribution and validation of Cx30 immunostaining

Using immunofluorescence, we first sought to characterize the general distribution of Cx30 in the ACC and validate antibody specificity. As expected^63,64^, Cx30-IR took the form of a very dense and clear punctate labeling which was highly enriched in cortical grey matter neuropil (**Fig.1A**) while virtually absent in adjacent white matter. Since Cx30 is known to be strongly associated with cerebral vasculature, where astrocytes establish numerous gap junctions at the perivascular endfeet^64–67^, we combined immunostainings for Cx30, vimentin, and GFAP to confirm that we could observe this distributional pattern. Vimentin was chosen because it is an intermediate filament expressed by cells in blood vessels walls as well as by a subpopulation of astrocytes^68^. The distribution of Cx30 was homogenous throughout cortical layers, and as illustrated in **Fig.1B**, Cx30 expression was particularly enriched along astrocyte endfeet lining blood vessels in highly vascularized areas. We further addressed the specificity of this distribution by immunostaining for GFAP together with both astrocyte-specific connexins, Cx30 and Cx43. As shown in **Fig.1C** and previously reported^45,65,67^, Cx30 and Cx43 highly colocalized at the level of GFAP+ endfeet along blood vessels, indicating that our Cx30 staining was reliable. We then quantified the overall density of Cx30 puncta in the ACC of controls and depressed suicides, segregating cases based on their known history of CA (**Fig.2A**). Surprisingly, although based on the literature we expected that Cx30 puncta density may be decreased in depressed suicides, we found no difference between groups (Kruskal-Wallis ANOVA: H(2,23) = 0.3268, P =0.84), regardless of the history of CA, as both depressed suicides with (DS-CA) and without (DS-nCA) history of CA showed Cx30 densities similar than controls. This was unlikely to be attributable to cofounding factors, as the density of Cx30 puncta was not correlated with age nor PMI (PMI vs Cx30 density: R^2^ = 0.01, P = 0.63; Age vs Cx30 density: R^2^ = 0.05, P = 0.33; **Fig.2D and E**). Because previous studies have linked antidepressant medication and alcohol abuse to changes in the expression of astrocytic connexins^52,69^, we compared Cx30 densities in cases with or without known history of substance abuse (**Fig.2B**), as well as cases with or without known antidepressant prescription during the last 3 months before death (**Figure 2C**). No significant effect of substance abuse (Kruskal-Wallis ANOVA: H(2,23) = 0.02247, P=0.99) or medication (Kruskal-Wallis ANOVA: H(2,23) = 0.3308, P=0.85) was found, supporting the notion that, at least in the ACC, overall Cx30 density is not altered as a function of psychopathology, medication or comorbid substance abuse.

**Figure 1.**
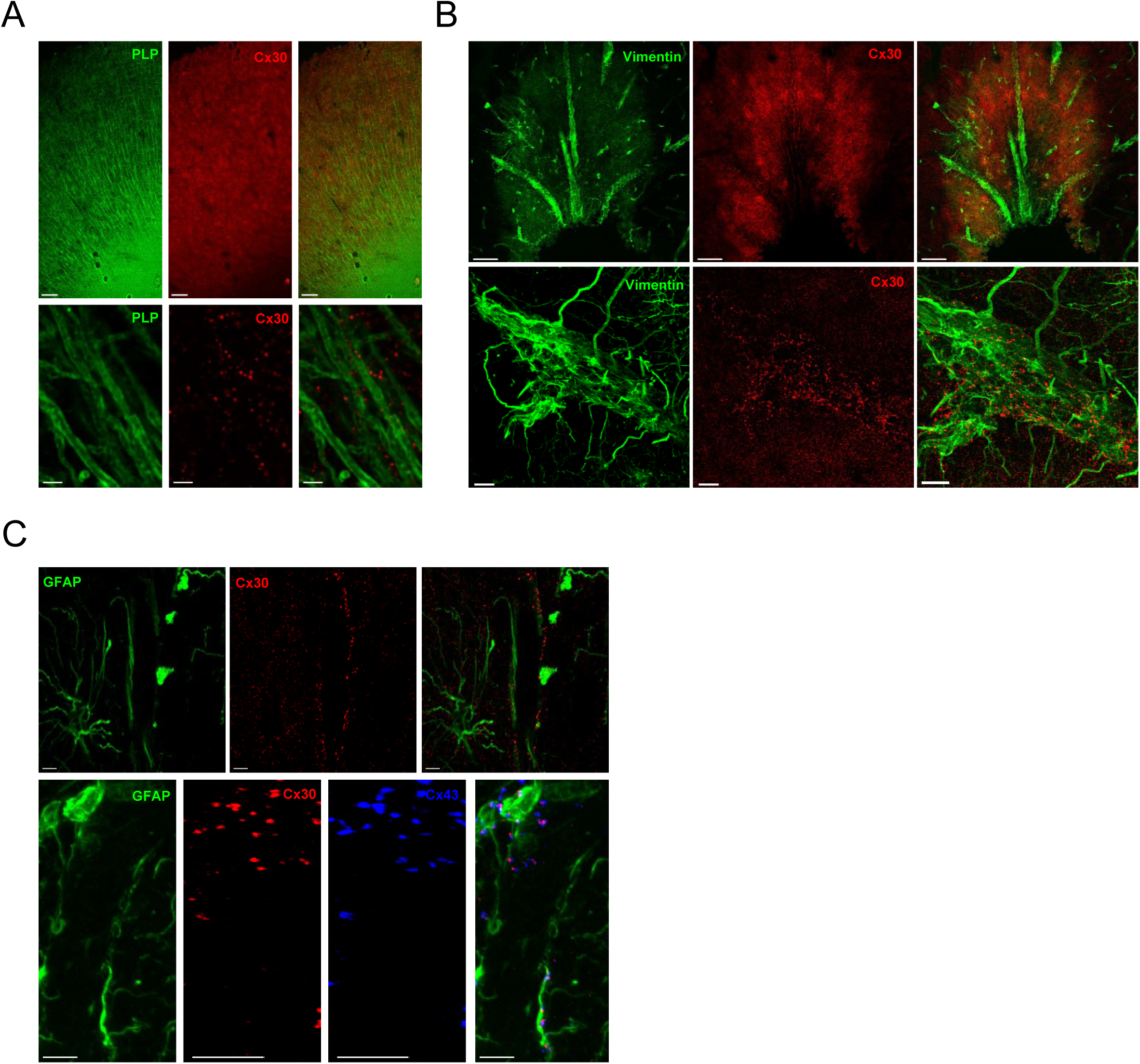
Distribution of connexin 30 immunoreactivity in the human anterior cingulate cortex. **(A)** Cx30 (red) at low (top panels) and high (bottom panels) magnification shows a strong level of expression in cortical grey matter but virtually no expression in deep white matter, with characteristic puncta-like immunoreactivity enriched in the neuropil. Myelin Proteolipid Protein (PLP) (green) was used to delineate white matter and visualize myelinated fibers. Cx30 surrounds and colocalizes with myelinated axons in cortical grey matter. Scale bars = 200 µm (top) and 2 µm (bottom). **(B)** Connexin 30 (red) is heavily enriched along brain vasculature and perivascular endfeet of astrocytes stained with vimentin (green). High magnification (bottom panels) shows strong colocalization of Cx30 puncta along blood vessels and perivascular endfeet, consistent with previous description of Cx30 distribution in the brain. Scale bars = 100 µm (top) and 10 µm (bottom) (C) Top panel shows the enrichment of Cx30-IR (red) along GFAP+ astrocytic endfeet (green) lining a blood vessel (BV). Bottom panel shows another BV surrounded by GFAP+ astrocytic fibers and perivascular endfeet (green), with strong colocalization of Cx30 and Cx43 (blue), as expected from the literature. Scale bars = 5 µm.

**Figure 2.**
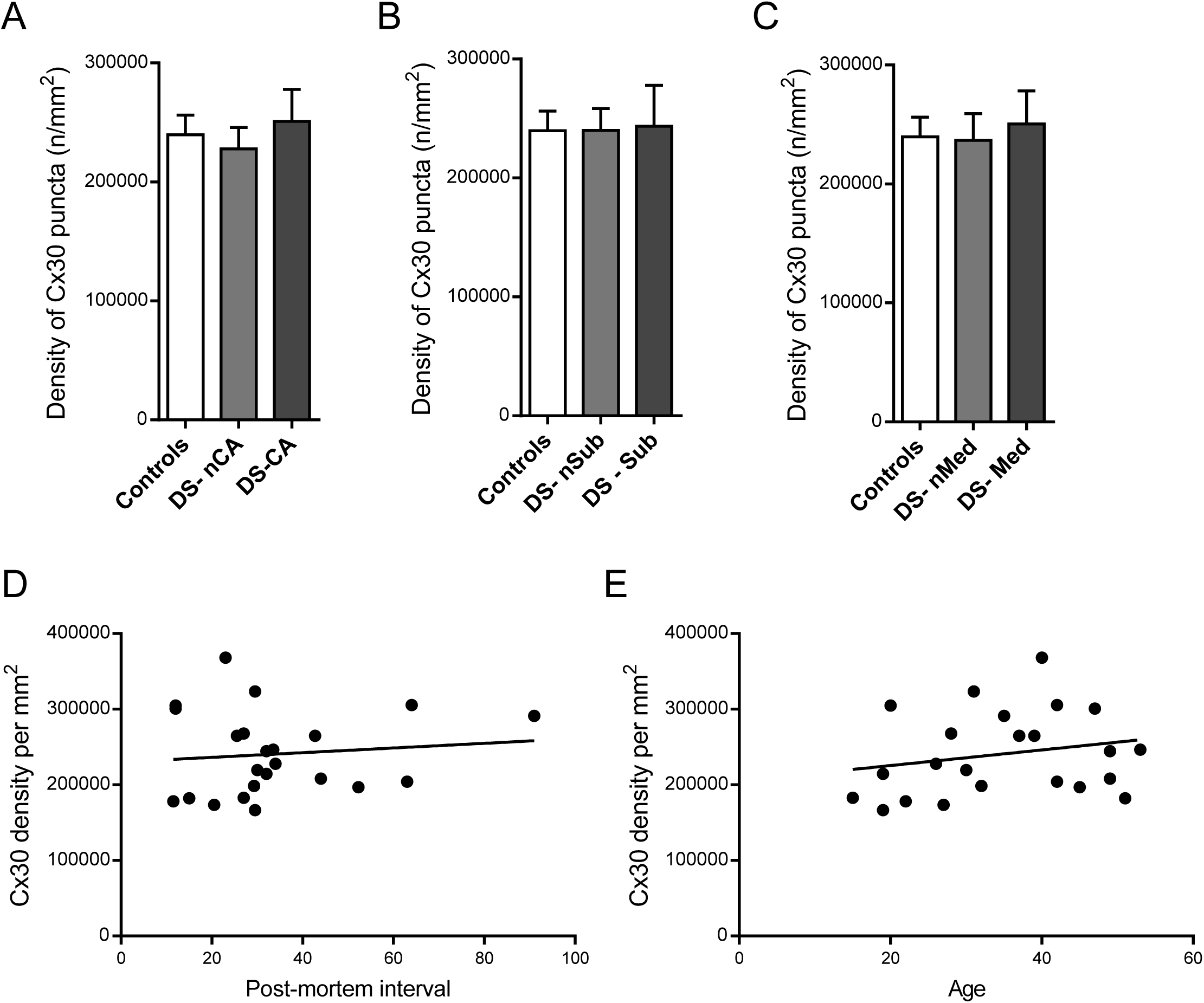
Depressed Suicides show no change in the density of Cx30-IR compared to controls in the anterior cingulate cortex. **(A)** Density of total Cx30-IR puncta in the ACC of depressed suicides without (DS-nCA, N=5)) or with (DS-CA, N=7) a history of child abuse and Controls (N=11). No difference between groups was found (Kruskal-Wallis ANOVA: H(2,23) = 0.3268, P =0.84). **(B)** No effect of substance abuse was found on the density of Cx30 in depressed suicides. Depressed suicides without (DS-nSub, N=7) or with (DS-Sub, N=5) a history of substance abuse showed similar densities of Cx30 compared to controls (N=11) (Kruskal-Wallis ANOVA: H(2,23) = 0.02247, P=0.99). **(C)** Previous medication with antidepressants had no effect on the densities of Cx30 in the ACC. Depressed suicides without (DS-nMed, N=8) or with (DS-Med), N=4) a known antidepressant prescription the last 3 months before death showed similar densities of Cx30 compared to controls (N=11) (Kruskal-Wallis ANOVA: H(2,23) = 0.3308, P=0.85). **(D)** No correlation between Cx30-IR and post-mortem interval (P=0.63) or **(E)** age (P = 0.33) was found, indicating overall that the absence of changes in Cx30 density between cases and controls is likely not linked to covariables.

### Decreased mapping of Cx30 onto oligodendrocyte cell bodies and myelinated fibers in depressed suicides

Because of the very high level of Cx30 expression and puncta labeling, small differences in Cx30 associated with specific cell-cell interactions could go unnoticed while assessing overall Cx30 distribution and density in the tissue. Since Cx30 is involved in astrocyte-oligodendrocyte gap junction coupling both at the soma and onto myelinated fibers^41^, we mapped Cx30-IR to oligodendrocytes and myelinated fibers to address possible changes specific to astrocyte-oligodendrocyte coupling (**Fig.3A**). Using Nogo-A-IR as a reliable marker of mature oligodendrocytes^19,70^, we first quantified the number of Cx30-IR puncta colocalizing with individual NogoA+ cells in layers 5/6 of the ACC (**Fig.3B**). Although the average mapping of Cx30 onto individual mature oligodendrocytes was relatively low, we found a significant difference between groups (Kruskal-Wallis ANOVA: H(2,23)= 9.09, P<0.05), with decreased Cx30-IR mapping to Nogo-A-IR cells in depressed suicides compared to controls, regardless of history of CA (Dunn’s multiple comparisons tests: CTRL vs DS-nCA and CTRL vs DS-CA: P<0.05). We then combined Myelin Proteolipid Protein (PLP) and Cx30 immunofluorescence to quantify the density of Cx30 mapping to myelinated fibers (**Fig.3C**). In accordance with decreased astrocyte-oligodendrocyte coupling at the soma, we also found differences between groups (Kruskal-Wallis ANOVA: H(2,24) = 7.471, P<0.05), with Cx30-IR decreased along PLP-IR myelinated axons in depressed suicides regardless of history of CA, although differences were more statistically robust in DS-CA (Dunn’s multiple comparison test, CTRL vs DS-CA: P<0.05), as only a trend was detectable between DS-nCA and controls (Dunn’s multiple comparison test, CTRL vs DS-nCA: P<0.1). Densities of Cx30 on PLP-IR myelinated axons were similar between DS-CA and DS-nCA (Dunn’s multiple comparison test, DS-nCA vs DS-CA: P=0.99).

**Figure 3.**
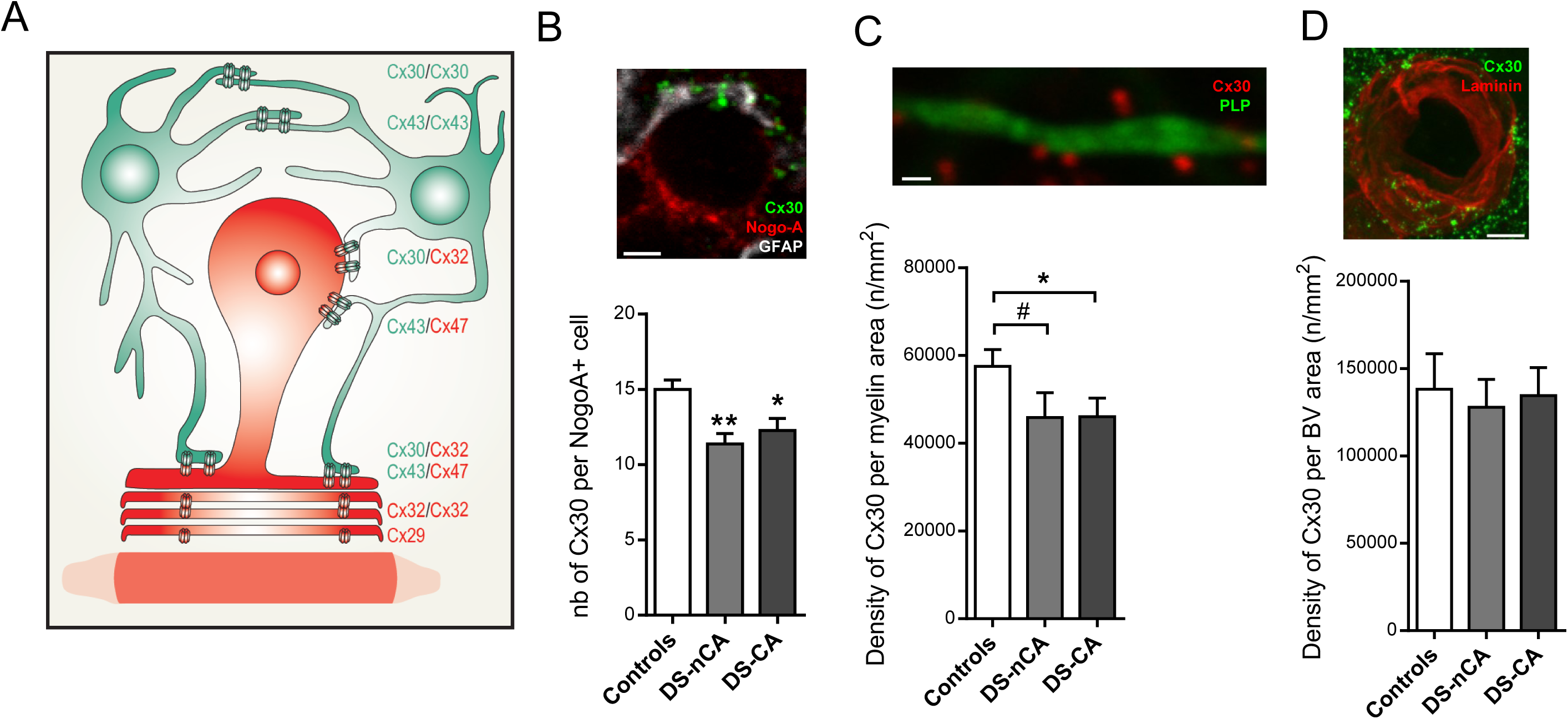
Specific decrease in Cx30-IR mapping to oligodendrocytes and myelinated fibers in the anterior cingulate of depressed suicides. **(A)** Drawing illustrating astrocyte-oligodendrocyte (A/O) gap junction coupling at the soma and around myelinated axons through heterotypic pairing of Cx30 or Cx43 on the astrocyte (colored in green) side, with Cx32 and Cx47 respectively on the oligodendrocyte (colored in red) side. Adapted from Orthmann-Murphy et al., 2007, *Journal of Neuroscience, 27, 13949–13957*. **(B)** Changes in A/O pairing between controls and depressed suicides was first addressed through quantification of Cx30-IR (green) mapping onto oligodendrocytes cells bodies stained with Nogo-A (red). Cx30-IR is found surrounding oligodendrocytes cells bodies and colocalizing in this example with GFAP+ astrocytes (grey). Depressed suicides, regardless of history of child abuse, showed a decrease in Cx30-IR mapping onto Nogo-A+ oligodendrocytes (Kruskal-Wallis ANOVA: H(2,23)= 9.09, P<0.05; Dunn’s multiple comparisons tests: Controls (N=11) vs DS-nCA (N=5) and Controls vs DS-CA (N=7), P<0.05). Scale bar = 2µm. **(C)** Changes in A/O pairing between controls and depressed suicides was also assessed though quantification of Cx30-IR (red) onto myelinated axons visualized with Myelin Proteolipid Protein (PLP, green). We found differences between groups (Kruskal-Wallis ANOVA: H(2,24) = 7.471, P<0.05), with Cx30-IR decreased along PLP-IR myelinated axons in depressed suicides regardless of history of CA, although differences were more statistically robust in DS-CA (Dunn’s multiple comparison test, Controls (N=10) vs DS-CA (N=11): P<0.05), as only a trend was detectable between DS-nCA and controls (Dunn’s multiple comparison test, Controls vs DS-nCA (N=6): P<0.1). Densities of Cx30 on PLP-IR myelinated axons were however similar between DS-CA and DS-nCA (Dunn’s multiple comparison test, DS-nCA vs DS-CA: P=0.99). Scale bar = 0.5 µm **(D)** Changes in Cx30-IR mapping seem specific to A/O gap junction coupling since no change between groups (Controls, N= 8; DS-nCA, N=8; DS-CA, N=9) was observed in Cx30-IR (green) mapping onto blood vessels stained with laminin (red) (Kruskal-Wallis ANOVA: H(2,25)= 0.2, P=0.9). *, P<0.05; **, P<01; #, 0.05<P<0.1 (versus Controls). Scale bar = 5 µm.

**Figure 4.**
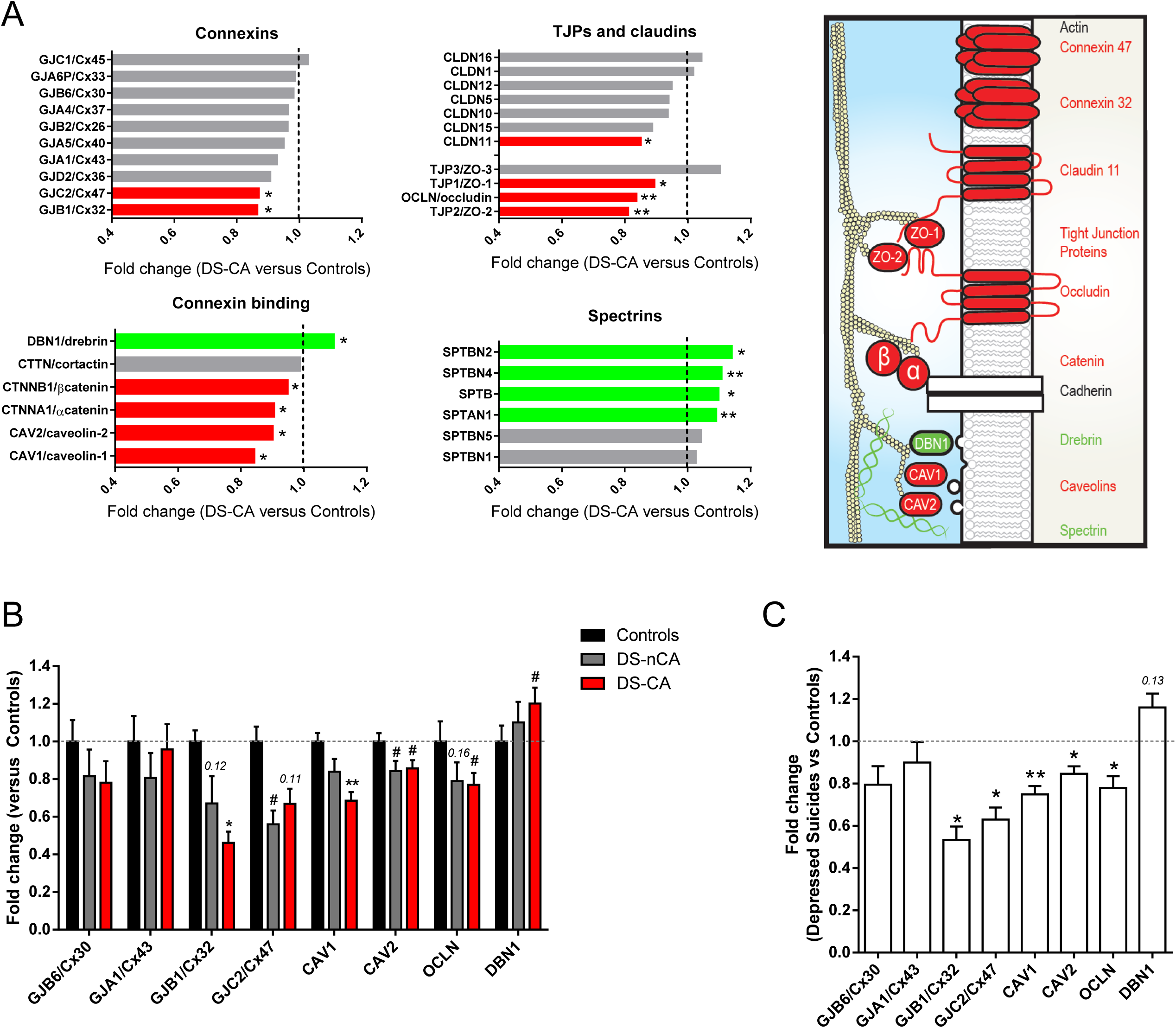
Genes involved in the scaffolding, trafficking and function of gap junction channels and cell-cell junctions are differentially expressed in the anterior cingulate cortex of depressed suicides. **(A)** RNA-sequencing results comparing samples from the anterior cingulate cortex of depressed suicides with a history of child abuse vs controls were screened to probe the differential expression of a literature-based list of genes related to gap junction channel scaffolding and function. We found that among genes coding for major connexins, depressed suicides showed a downregulation specifically for the two oligodendrocyte-specific connexins Cx32 (GJB1) and Cx47 (GJC2). Major tight junction proteins (TJPs) such as occluding (OCLN), zona occludens-1 and −2 (ZO1 and ZO2), as well as claudin-11/OSP (CLDN11), an essential component of myelin interlamellar tight junctions were also found to be downregulated in depressed suicides. Genes coding for connexins binding partners and regulators of gap junction mediated intercellular communication, such as caveolin-1 and −2 (CAV1 and CAV2) and alpha/beta catenins (CTNNA1 and CTNNB1) were also found to be downregulated. Of note, several genes coding for actin-binding proteins thought to bridge connexins to actin microfilament cytoskeleton, such as drebrin (developmentally regulated brain protein, DBN1) and spectrins were found to be collectively upregulated in depressed suicides compared to controls. Red and green bars indicate significant downregulation and upregulation respectively, while grey bars indicate no significant change in gene expression. The drawing on the right depicts the suggested interactions between these different components of the gap junction nexus and connexin-interacting proteins, adapted from Dbouk et al., *Cell Commun Signal 2009;* ***7****: 4*. Red and green indicate components found to be downregulated or upregulated, respectively, in the RNA-sequencing dataset. **(B)** Because the RNA-sequencing dataset was generated comparing samples from controls and depressed suicides with a history of child abuse (DS-CA), a subset of these differentially expressed genes was selected for validation with quantitative RT-PCR and adding a third group of depressed suicides with no history of child abuse (DS-nCA). Globally and consistent with the RNA-sequencing results, DS-CA subjects showed a downregulation of Cx32/GJB1, Cx47/GJC2, CAV1 and CAV2, OCLN and an upregulation of DBN1 (Controls (N=16-23) vs DS-CA (N=19-25), *=P<0.05, **=P<0.01, #=P<0.1), although these results were not as statistically robust as observed in the RNA-sequencing dataset. Importantly these changes did not seem specific to a history of child abuse, as no significant difference was found between DS-CA and DS-nCA for any of these genes, while DS-nCA showed similar trends towards decreased expression of Cx32/GJB1, Cx47/GJC2, CAV2 and OCLN (Controls (N=16-23) vs DS-nCA (N=12-24), #=P<0.1). **(C)** Since changes in gene expression were not found to be specific to a history of child abuse, we combined both depressed suicides groups to increase our statistical power and to highlight significant downregulations of oligodendrocyte connexins Cx32/GJB1 and Cx47/GJC2, CAV1 and CAV2 as well as OCLN (Controls (N=16-23) vs Depressed Suicides (N=33-48), *=P<0.05, **=P<0.01). The upregulation of DBN1 observed in the RNA-seq dataset was however not validated, although a small trend towards increased expression was detectable (Controls (N=23) vs Depressed Suicides (N=36), P=0.13).

To address whether changes in Cx30 distribution could also be observed between astrocytes and other cell types, we used a similar approach to assess the Cx30-IR associated with cerebral vasculature immunolabeled for laminin (**Fig.3D**). Unlike what was observed on Nogo-A-IR cells and PLP-IR axons, the density of Cx30-IR along laminin-IR blood vessels was similar between groups (Kruskal-Wallis ANOVA: H(2,25)= 0.2, P=0.9). These results highlight that changes in Cx30-mediated gap junction coupling in the ACC of depressed suicides may be specific to astrocyte-oligodendrocyte communication.

### Downregulation of oligodendrocyte-specific connexins and scaffolding components of gap junction channels in the ACC of depressed suicides

Changes in gap junction coupling are likely to involve transcriptomic alterations of genes essential for the synthesis of GJC components, their scaffolding, trafficking and function. To complement our histological approach, we screened the gene expression of some key regulators of the gap junction proteome from a previously published RNA-sequencing dataset comparing controls and matched depressed suicides in the ACC^18^. This dataset had allowed to find a global downregulation of oligodendrocyte-related genes, as well as morphometric impairments of myelin in depressed suicides with a history of CA. Here, a non-exhaustive literature-based list of genes was screened that included major connexins and some of their known binding partners, such as cytoskeletal and scaffolding components of the GJC (**Fig.4A and B**). The latter included zona occludens 1 (ZO-1), ZO-2 and ZO-3, spectrin, drebrin and catenins, which together form a network of proteins that bridges connexins to microfilaments and microtubules and interact with occludins, claudins and cadherins to form junctional nexuses^71–74^.

Among connexin genes, only those coding for two oligodendrocyte-specific connexins, Cx32 (GJB1) and Cx47 (GJC2), were found to be downregulated in depressed suicides (P<0.05). These connexins are binding partners of the astrocyte-specific connexins Cx30 and Cx43, respectively, which pair together to form functional GJCs between astrocytes and oligodendrocytes^41,43^. Moreover, two major connexin-interacting proteins, alpha- and beta-catenins^75,76^, both adherens junction components, were also downregulated in cases versus controls (P<0.05). Several major cytoskeletal components essential for the scaffolding of GJCs and junctional nexus were also found to be differentially expressed. The major tight junction proteins ZO-1 and ZO-2, interacting with each other and with connexins to regulate gap junction plaque size and distribution^71,77,78^, were downregulated in samples from depressed suicides (P<0.05 and P<0.01 respectively). We also screened for other core proteins of tight junctions, such as occludin and claudins, which are also part of the junctional nexus, some of which co-immunoprecipitate with the oligodendrocyte-expressed Cx32^79,80^. Again, occludin was significantly downregulated in depressed suicides (P<0.01), while among the claudin family of proteins, the only member found to be differentially expressed between groups was CLDN11 (claudin-11, P<0.05), an essential component of myelin interlamellar tight junctions^81–83^, also suggesting that cell-cell junctions involving oligodendrocytes may be particularly impaired in depressed suicides.

Caveolin-1 and Caveolin-2, two major regulators of connexin trafficking and assembly, as well as gap junction mediated intercellular communication^84,85^, were also found to be downregulated in depressed suicides (P<0.05). Finally, of the key actin-binding proteins important to bridge connexins to actin microfilament cytoskeleton^86–90^, DBN1 (developmentally regulated brain protein, drebrin, P<0.05), SPTAN1 (Alpha-II Spectrin, P<0.01), SPTB (beta-I spectrin, P<0.05), SPTBN2 (beta-III spectrin, P<0.05), and SPTBN4 (Beta-IV Spectrin, P<0.01) were all upregulated in depressed suicides compared to controls.

A subset of these differentially expressed genes was selected for validation with quantitative RT-PCR. We focused on the connexins enriched in astrocytes (Cx30 and cx43) and oligodendrocytes (Cx32 and Cx47) as well as CAV1, CAV2, OCLN and DBN1 (**Fig.4C and D**). Importantly, because in our RNA-sequencing study all depressed suicides had a history of CA, which may specifically lead to oligodendrocyte abnormalities, qPCR validation was performed in both DS-nCA and DS-CA groups similarly to our histological experiments, to clarify whether observed changes were associated with psychopathology or were specific to history of CA. Overall, regardless of CA, depressed suicides displayed a general downregulation of Cx32, Cx47, CAV1 and CAV2 as well as OCLN expression and an upregulation of DBN1 expression compared to controls (**Fig.4C**). These changes were however not as statistically robust as previously observed using RNA-seq. Importantly, because these changes did not seem specific to a history of CA, as no significant difference was found between DS-CA and DS-nCA for any of these genes, we combined both depressed suicides groups (**Fig4D**) and found significant downregulations of oligodendrocyte connexins Cx32 and Cx47, CAV1 and CAV2 as well as OCLN (CTRL vs Depressed Suicides, P<0.05). The upregulation of DBN1 observed in the RNA-seq dataset was however not validated, although a small trend towards increased expression was detectable (CTRL vs Depressed Suicides, P=0.13).

## DISCUSSION

In this study, we aimed to address the hypothesis that cellular communication between astrocytes and oligodendrocytes could potentially be impaired in depressed suicides, as previous evidence had linked changes in the expression of astrocytic connexins as well as myelin and oligodendrocyte function impairments in this population. We characterized the distribution of Cx30 in the human post-mortem ACC by immunolabeling, and found that Cx30-IR is selectively decreased in somas of mature oligodendrocytes and along myelinated fibers in depressed suicides, suggesting that Cx30-mediated gap junction coupling between astrocytes and oligodendrocytes may be disrupted. This was supported by the reduced expression of genes coding for the two major oligodendrocyte-specific connexins, Cx32 and Cx47, which upon pairing with the astrocytic connexins Cx30 and Cx43 allow the formation of functional GJCs between these two glial cell types. Additional key components of the gap junction proteome and cell-cell junctions, in particular components essential for the scaffolding, trafficking and function of GJCs, were also found to be downregulated in depressed suicides. Altogether, these results highlight that glial cell communication through cell-cell junctions may be an important feature of the pathophysiology of depression, possibly bridging the cumulative evidence of astrocytes and oligodendrocytes abnormalities previously observed in this population.

Several post-mortem studies, including from our group, have consistently reported a downregulation in the expression of astrocytic connexins in frontal regions of suicides. Ernst et al. first found both Cx30 and Cx43 to be strongly downregulated in the dorsolateral prefrontal cortex of suicides^91^. Nagy et al. further showed that this phenomenon also occurred in other cortical and subcortical brain regions, namely the prefrontal, primary motor and visual cortex, the dorsomedial thalamic nucleus, and the caudate nucleus^7^. Finally, Miguel-Hidalgo and collaborators also reported that Cx43 expression was decreased in the orbitofrontal cortex of depressed individuals versus controls^52^. We were therefore surprised that this result was not replicated in our cohort of subjects, perhaps highlighting a regional specificity of the ACC with regards to astrocytic changes associated with psychopathology. This discrepancy was not driven by covariates, history of substance dependence or antidepressant treatment. In agreement with our results for Cx30 and Cx43 in the ACC, other canonical markers of astrocytes previously found to be downregulated in BA8/9 of depressed suicides, alongside astrocytic connexins, were not affected in the ACC. These markers include ALDH1L1, GLUL, and Sox9, an astrocyte-enriched transcription factor previously shown to bind to and potently regulate Cx30 expression^91^. Considering that astrocyte connexins are engaged in a wide range of GJCs between astrocytes and other cell types, we hypothesize that astrocytes dysfunction may not be a critical pathophysiological feature in the ACC, but that their interaction with other cell types may be still be compromised through changes occurring in some of their interacting partners. In that scenario whole levels of Cx30 expression may not be indicative of discrete changes occurring in other cell types. We were particularly interested in glial communication between astrocytes and oligodendrocytes, as we previously showed that cortical myelination is critically impaired in the ACC of depressed suicides^18^, and that A/O GJCs are essential for the maintenance of myelin and oligodendrocyte function^40^.

Our results indicated that indeed, while only representing a small fraction of the Cx30 localization, Cx30 mapping to mature oligodendrocytes and myelinating fibers, but not cerebral vasculature, is selectively decreased in depressed suicides. In accordance with these immunohistological results, we found that the gene expression of the canonical partner of Cx30 on the oligodendrocyte side, Cx32, was also significantly downregulated. Taken together, these observations strengthen our interpretation that A/O gap junction coupling may be impaired in depression, possibly through changes occurring in oligodendrocytes. Interestingly, the other oligodendrocyte-specific connexin, Cx47, which associates with Cx43, was also downregulated in depressed suicides. Although we did not quantify the mapping of Cx43 onto oligodendrocytes and myelinated fibers because of antibody incompatibilities, these results also suggest that A/O gap junction coupling impairment may not be specific to Cx30-mediated GJCs but likely extend to Cx43-Cx47 pairing as well. While we focused on A/O coupling, it would be interesting to use this approach to explore other cell-cell interactions, in particular O/O coupling through homotypic pairing of Cx32 and Cx47, both of which are downregulated in the ACC of depressed suicides.

To our knowledge, mapping connexin expression in situ onto different cell types has never been achieved previously in post-mortem brain tissue, in which immunofluorescence can be challenging to perform. This strategy has some limitations, however. Given the limited resolution of confocal microscopy, it is impossible to infer whether the decreased mapping of Cx30 to oligodendrocytes and myelinated fibers represents a loss of Cx30 pairing to its partner Cx32 or changes in hemichannel trafficking. While a more direct approach consisting in mapping the colocalization of Cx30 with Cx32 onto oligodendrocytes or their interaction by co-immunoprecipitation would have certainly allowed a more precise assessment of A/O coupling, this proved challenging in our hands because of antibodies incompatibility. Despite this limitation, our immunohistological approach was supported by strong evidence showing impaired expression of key components of the gap junction proteome in the ACC of depressed suicides. Alongside reduced expression of the two oligodendrocyte connexins Cx32 and Cx47, major cytoskeletal scaffolding components and regulators of gap junction trafficking and assembly showed a collective downregulation. While some of those are major tight and adherens junction proteins, such as occludin, ZO1 and ZO2, or claudin-11, strong evidence highlights their role in supporting the scaffolding and function GJCs^71,73,86^. Alongside our observed changes in actin-binding proteins bridging connexins to cytoskeletal microfilaments, such as spectrins and catenins, this hints towards a convergent disruption of the structural integrity of cell-cell junctions. In view of our immunohistological results and reduced expression of Cx32 and Cx47, we hypothesize that this may reflect disruptions of gap junction integrity, although it is likely that such global effects may translate in other cell-cell junctions, in particular tight and adherens junctions. Tight junctions make up the radial component of CNS myelin, acting as an interlaminar diffusion barrier and structural support^92^. The fact that claudin-11, the major component of oligodendrocyte tight junctions^82^, was the only member of the claudin family to be differently expressed in depressed suicides suggests that transcriptional changes associated with psychopathology are particularly prominent in OLs.

What upstream causes and mechanisms could underlie the disrupted gap junction connectivity between astrocytes and oligodendrocytes in the ACC of depressed suicides? One possible answer could be neuroinflammation, which has been increasingly associated with depression and suicide^93–95^. Indeed, several pro-inflammatory molecules are thought to alter the composition and structure of tight junctions by modulating some of their key components such as occludin, claudins, and zona occludens proteins^96–98^, thereby affecting their permeability and function. This has been documented in the CNS, and disruption of blood brain barrier integrity through alterations in tight junctions has recently emerged as an interesting pathophysiological feature of depression^99^. With regards to connexins and gap junction integrity, chronic inflammation, microglial activation and astrogliosis have also been associated with Cx32 and Cx47 downregulation, and widespread loss of O/O and O/A gap junctions^100–104^. Notably, loss of Cx32 GJs at oligodendrocyte cells bodies and along myelinated fibers in grey matter were previously reported in multiple sclerosis^103^, similar to our observations showing decreased Cx30-IR on oligodendrocyte cell bodies and myelinated axons. Given that increased microglia/macrophage activity has been reported in the ACC of depressed suicides^94^, the present evidence of cell-cell junctions and altered A/O communication may represent a consequence of depression-associated low-level chronic neuroinflammation.

We recently reported a robust downregulation of the oligodendrocyte transcriptome and decreased myelination of small caliber axons in the ACC of depressed suicides^18^. Remarkably, these effects were associated with a history of severe CA, but were absent in a matched sample of depressed suicides without history of CA. The present study provides evidence that altered A/O communication in the ACC may be a common neurobiological feature of major depression, irrespective of history of early-life adversity, and could possibly be linked to the characteristic changes in ACC volume and function described in depressed patients^31,105^. It can be further hypothesized that the more global impact of CA on oligodendrocytes and myelination may predispose to these changes and mediate specific alterations of inter-regional connectivity found in depressed individuals exposed to early-life adversity^106^.

In conclusion, our study suggests that the astrocyte-oligodendrocyte gap junction connectivity is decreased in the ACC of depressed suicides, possibly through decreased Cx30 coupling to oligodendrocyte- and myelin-specific connexins. This is associated with a convergent downregulation of key structural components and regulators of cell-cell junctions. Because A/O GJCs play an important role in providing metabolic support to oligodendrocyte-lineage cells, regulate precursor cells proliferation and differentiation, and are essential for myelin integrity, we argue that our findings may relate to reported alterations of myelination previously observed in depressed suicides and further support glial dysfunction as a major hallmark of depression. Ultimately, as suggested by recent preclinical studies^54,69,107^, targeting connexins and cell-cell junctions could prove useful in the search of novel therapeutic approaches for depression.

## ACKNOWLEDGMENTS

This research was funded by grants from CIHR (PJT-156346) and the AFSP (SRG-0-088-15) to N.M. A.T. was supported by a TD postdoctoral fellowship, and currently by an AFSP Young Investigator Innovation Grant (YIG-0-146-17). P-E.L. was supported by scholarships from the American Foundation of Suicide Prevention, the Fondation Deniker, the Fondation pour la Recherche Médicale, and the UNAFAM (‘Union nationale de familles et amis de personnes malades et/ou handicapées psychiques’). The authors wish to thank the Douglas-Brain Canada Brain Bank (DBCBB) staff as well as the Douglas Institute’s Molecular and Cellular Microscopy Platform coordinator for their precious assistance. Both of these research platforms are funded by a Core Facilities and Technology Development grant from Healthy Brains for Healthy Lives (CFREF). The DBCBB is further supported by the RQSHA (FRQ-S).

## REFERENCES

1. Miguel-Hidalgo JJ, Baucom C, Dilley G, Overholser JC, Meltzer HY, Stockmeier CA et al. Glial fibrillary acidic protein immunoreactivity in the prefrontal cortex distinguishes younger from older adults in major depressive disorder. Biol Psychiatry 2000; 48: 861–73.

2. Si X, Miguel-Hidalgo JJ, O’Dwyer G, Stockmeier CA, Rajkowska G. Age-Dependent Reductions in the Level of Glial Fibrillary Acidic Protein in the Prefrontal Cortex in Major Depression. Neuropsychopharmacology 2004; 29: 2088–2096.

3. Rajkowska G, Hughes J, Stockmeier CA, Javier Miguel-Hidalgo J, Maciag D. Coverage of Blood Vessels by Astrocytic Endfeet Is Reduced in Major Depressive Disorder. Biol Psychiatry 2013; 73: 613–621.

4. Gittins RA, Harrison PJ. A morphometric study of glia and neurons in the anterior cingulate cortex in mood disorder. J Affect Disord 2011; 133: 328–332.

5. Torres-Platas SG, Hercher C, Davoli MA, Maussion G, Labonté B, Turecki G et al. Astrocytic hypertrophy in anterior cingulate white matter of depressed suicides. Neuropsychopharmacology 2011; 36: 2650–8.

6. Torres-Platas SG, Nagy C, Wakid M, Turecki G, Mechawar N. Glial fibrillary acidic protein is differentially expressed across cortical and subcortical regions in healthy brains and downregulated in the thalamus and caudate nucleus of depressed suicides. Mol Psychiatry 2016; 21: 509–515.

7. Nagy C, Torres-Platas SG, Mechawar N, Turecki G. Repression of Astrocytic Connexins in Cortical and Subcortical Brain Regions and Prefrontal Enrichment of H3K9me3 in Depression and Suicide. Int J Neuropsychopharmacol 2016; 20: pyw071.

8. Nagy C, Suderman M, Yang J, Szyf M, Mechawar N, Ernst C et al. Astrocytic abnormalities and global DNA methylation patterns in depression and suicide. Mol Psychiatry 2015; 20: 320–8.

9. Rajkowska G, Legutko B, Moulana M, Syed M, Romero DG, Stockmeier CA et al. Astrocyte pathology in the ventral prefrontal white matter in depression. J Psychiatr Res 2018; 102: 150–158.

10. Czéh B, Nagy SA. Clinical Findings Documenting Cellular and Molecular Abnormalities of Glia in Depressive Disorders. Front Mol Neurosci 2018; 11: 56.

11. Leng L, Zhuang K, Liu Z, Huang C, Gao Y, Chen G et al. Menin Deficiency Leads to Depressive-like Behaviors in Mice by Modulating Astrocyte-Mediated Neuroinflammation. Neuron 2018; 100: 551–563.e7.

12. Banasr M, Duman RS. Glial Loss in the Prefrontal Cortex Is Sufficient to Induce Depressive-like Behaviors. Biol Psychiatry 2008; 64: 863–870.

13. Almeida RG, Lyons DA. On Myelinated Axon Plasticity and Neuronal Circuit Formation and Function. J Neurosci 2017; 37: 10023–10034.

14. Gibson EM, Geraghty AC, Monje M. Bad wrap: Myelin and myelin plasticity in health and disease. Dev Neurobiol 2018; 78: 123–135.

15. Forbes TA, Gallo V. All Wrapped Up: Environmental Effects on Myelination. Trends Neurosci 2017; 40: 572–587.

16. Uranova NA, Vostrikov VM, Orlovskaya DD, Rachmanova VI. Oligodendroglial density in the prefrontal cortex in schizophrenia and mood disorders: a study from the Stanley Neuropathology Consortium. Schizophr Res 2004; 67: 269–75.

17. Uranova N, Orlovskaya D, Vikhreva O, Zimina I, Kolomeets N, Vostrikov V et al. Electron microscopy of oligodendroglia in severe mental illness. Brain Res Bull 2001; 55: 597–610.

18. Lutz P-E, Tanti A, Gasecka A, Barnett-Burns S, Kim JJ, Zhou Y et al. Association of a History of Child Abuse With Impaired Myelination in the Anterior Cingulate Cortex: Convergent Epigenetic, Transcriptional, and Morphological Evidence. Am J Psychiatry 2017;: appiajp201716111286.

19. Tanti A, Kim JJ, Wakid M, Davoli M-A, Turecki G, Mechawar N. Child abuse associates with an imbalance of oligodendrocyte-lineage cells in ventromedial prefrontal white matter. Mol Psychiatry 2018; 23: 2018–2028.

20. Hayashi Y, Nihonmatsu-Kikuchi N, Yu X, Ishimoto K, Hisanaga S, Tatebayashi Y. A novel, rapid, quantitative cell-counting method reveals oligodendroglial reduction in the frontopolar cortex in major depressive disorder. Mol Psychiatry 2011; 16: 1156–1158.

21. Rajkowska G, Mahajan G, Maciag D, Sathyanesan M, Iyo AH, Moulana M et al. Oligodendrocyte morphometry and expression of myelin - Related mRNA in ventral prefrontal white matter in major depressive disorder. J Psychiatr Res 2015; 65: 53–62.

22. Edgar N, Sibille E. A putative functional role for oligodendrocytes in mood regulation. Transl Psychiatry 2012; 2: e109.

23. Aston C, Jiang L, Sokolov BP. Transcriptional profiling reveals evidence for signaling and oligodendroglial abnormalities in the temporal cortex from patients with major depressive disorder. Mol Psychiatry 2005; 10: 309–322.

24. Kim S, Webster MJ. Correlation analysis between genome-wide expression profiles and cytoarchitectural abnormalities in the prefrontal cortex of psychiatric disorders. Mol Psychiatry 2010; 15: 326–336.

25. Laine MA, Trontti K, Misiewicz Z, Sokolowska E, Kulesskaya N, Heikkinen A et al. Genetic Control of Myelin Plasticity after Chronic Psychosocial Stress. eneuro 2018; 5: ENEURO.0166-18.2018.

26. Gao Y, Ma J, Tang J, Liang X, Huang C-X, Wang S et al. White matter atrophy and myelinated fiber disruption in a rat model of depression. J Comp Neurol 2017; 525: 1922–1933.

27. Miyata S, Taniguchi M, Koyama Y, Shimizu S, Tanaka T, Yasuno F et al. Association between chronic stress-induced structural abnormalities in Ranvier nodes and reduced oligodendrocyte activity in major depression. Sci Rep 2016; 6: 23084.

28. Tham MW, Woon PS, Sum MY, Lee T-S, Sim K. White matter abnormalities in major depression: evidence from post-mortem, neuroimaging and genetic studies. J Affect Disord 2011; 132: 26–36.

29. Kim B, Oh J, Kim M-K, Lee S, Suk Tae W, Mo Kim C et al. White matter alterations are associated with suicide attempt in patients with panic disorder. J Affect Disord 2015; 175C: 139–146.

30. Peng H, Ning Y, Zhang Y, Yang H, Zhang L, He Z et al. White-matter density abnormalities in depressive patients with and without childhood neglect: A voxel-based morphometry (VBM) analysis. Neurosci Lett 2013; 550: 23–28.

31. Drevets WC, Price JL, Furey ML. Brain structural and functional abnormalities in mood disorders: implications for neurocircuitry models of depression. Brain Struct Funct 2008; 213: 93–118.

32. Johansen-Berg H, Gutman DA, Behrens TEJ, Matthews PM, Rushworth MFS, Katz E et al. Anatomical connectivity of the subgenual cingulate region targeted with deep brain stimulation for treatment-resistant depression. Cereb Cortex 2008; 18: 1374–1383.

33. Steffens DC, Taylor WD, Denny KL, Bergman SR, Wang L. Structural Integrity of the Uncinate Fasciculus and Resting State Functional Connectivity of the Ventral Prefrontal Cortex in Late Life Depression. PLoS One 2011; 6: e22697.

34. Birn RM, Shackman a J, Oler J a, Williams LE, McFarlin DR, Rogers GM et al. Evolutionarily conserved prefrontal-amygdalar dysfunction in early-life anxiety. Mol Psychiatry 2014; 19: 915–922.

35. Koolschijn PCMP, van Haren NEM, Lensvelt-Mulders GJLM, Hulshoff Pol HE, Kahn RS. Brain volume abnormalities in major depressive disorder: A meta-analysis of magnetic resonance imaging studies. Hum Brain Mapp 2009; 30: 3719–3735.

36. Domingues HS, Portugal CC, Socodato R, Relvas JB. Oligodendrocyte, Astrocyte, and Microglia Crosstalk in Myelin Development, Damage, and Repair. Front cell Dev Biol 2016; 4: 71.

37. Dutta DJ, Woo DH, Lee PR, Pajevic S, Bukalo O, Huffman WC et al. Regulation of myelin structure and conduction velocity by perinodal astrocytes. Proc Natl Acad Sci U S A 2018; 115: 11832–11837.

38. Kıray H, Lindsay SL, Hosseinzadeh S, Barnett SC. The multifaceted role of astrocytes in regulating myelination. Exp Neurol 2016; 283: 541–9.

39. Li J, Zhang L, Chu Y, Namaka M, Deng B, Kong J et al. Astrocytes in Oligodendrocyte Lineage Development and White Matter Pathology. Front Cell Neurosci 2016; 10: 119.

40. Nualart-Marti A, Solsona C, Fields RD. Gap junction communication in myelinating glia. Biochim Biophys Acta - Biomembr 2013; 1828: 69–78.

41. Orthmann-murphy JL, Abrams CK, Scherer SS. Gap Junctions Couple Astrocytes and Oligodendrocytes. 2008. doi:10.1007/s12031-007-9027-5.

42. Evans WH, Martin PEM. Gap junctions: structure and function (Review). Mol Membr Biol 2002; 19: 121–136.

43. Magnotti LM, Goodenough DA, Paul DL. Functional heterotypic interactions between astrocyte and oligodendrocyte connexins. Glia 2011; 59: 26–34.

44. Maglione M, Tress O, Haas B, Karram K, Trotter J, Willecke K et al. Oligodendrocytes in mouse corpus callosum are coupled via gap junction channels formed by connexin47 and connexin32. Glia 2010; 58: 1104–1117.

45. Nagy JI, Rash JE. Connexins and gap junctions of astrocytes and oligodendrocytes in the CNS. Brain Res Brain Res Rev 2000; 32: 29–44.

46. Rash JE, Yasumura T, Davidson KG, Furman CS, Dudek FE, Nagy JI. Identification of cells expressing Cx43, Cx30, Cx26, Cx32 and Cx36 in gap junctions of rat brain and spinal cord. Cell Commun Adhes 2001; 8: 315–20.

47. Orthmann-Murphy JL, Freidin M, Fischer E, Scherer SS, Abrams CK. Two Distinct Heterotypic Channels Mediate Gap Junction Coupling between Astrocyte and Oligodendrocyte Connexins. J Neurosci 2007; 27: 13949–13957.

48. Menichella DM, Goodenough DA, Sirkowski E, Scherer SS, Paul DL. Connexins are critical for normal myelination in the CNS. J Neurosci 2003; 23: 5963–73.

49. Lutz SE, Zhao Y, Gulinello M, Lee SC, Raine CS, Brosnan CF. Deletion of Astrocyte Connexins 43 and 30 Leads to a Dysmyelinating Phenotype and Hippocampal CA1 Vacuolation. J Neurosci 2009; 29: 7743–7752.

50. May D, Tress O, Seifert G, Willecke K. Connexin47 protein phosphorylation and stability in oligodendrocytes depend on expression of Connexin43 protein in astrocytes. J Neurosci 2013; 33: 7985–96.

51. Tress O, Maglione M, May D, Pivneva T, Richter N, Seyfarth J et al. Panglial Gap Junctional Communication is Essential for Maintenance of Myelin in the CNS. J Neurosci 2012; 32: 7499–7518.

52. Miguel-Hidalgo JJ, Wilson BA, Hussain S, Meshram A, Rajkowska G, Stockmeier CA. Reduced connexin 43 immunolabeling in the orbitofrontal cortex in alcohol dependence and depression. J Psychiatr Res 2014; 55: 101–9.

53. Miguel-hidalgo J, Moulana M, Deloach PH, Rajkowska G. Chronic Unpredictable Stress Reduces Immunostaining for Connexins 43 and 30 and Myelin Basic Protein in the Rat Prelimbic and Orbitofrontal Cortices. 2018. doi:10.1177/2470547018814186.

54. Sun J-D, Liu Y, Yuan Y-H, Li J, Chen N-H. Gap junction dysfunction in the prefrontal cortex induces depressive-like behaviors in rats. Neuropsychopharmacology 2012; 37: 1305–20.

55. Pannasch U, Rouach N. Emerging role for astroglial networks in information processing: from synapse to behavior. Trends Neurosci 2013; 36: 405–417.

56. Dossi E, Vasile F, Rouach N. Human astrocytes in the diseased brain. Brain Res Bull 2018; 136: 139–156.

57. Rial D, Lemos C, Pinheiro H, Duarte JM, Gonçalves FQ, Real JI et al. Depression as a Glial-Based Synaptic Dysfunction. Front Cell Neurosci 2016; 9: 521.

58. Teicher MH, Samson JA. Childhood maltreatment and psychopathology: A case for ecophenotypic variants as clinically and neurobiologically distinct subtypes. Am J Psychiatry 2013; 170: 1114–33.

59. Dumais A, Lesage AD, Alda M, Rouleau G, Dumont M, Chawky N et al. Risk factors for suicide completion in major depression: a case-control study of impulsive and aggressive behaviors in men. Am J Psychiatry 2005; 162: 2116–24.

60. Bifulco A, Brown GW, Harris TO. Childhood Experience of Care and Abuse (CECA): a retrospective interview measure. J Child Psychol Psychiatry 1994; 35: 1419–35.

61. Bifulco A, Brown GW, Lillie A, Jarvis J. Memories of childhood neglect and abuse: corroboration in a series of sisters. J Child Psychol Psychiatry 1997; 38: 365–74.

62. Mai JK, Paxinos G, Voss T. Atlas of the human brain. Elsevier Acad. Press, 2008.

63. Kunzelmann P, Schröder W, Traub O, Steinhäuser C, Dermietzel R, Willecke K. Late onset and increasing expression of the gap junction protein connexin30 in adult murine brain and long-term cultured astrocytes. Glia 1999; 25: 111–9.

64. Nagy JI, Patel D, Ochalski PA, Stelmack GL. Connexin30 in rodent, cat and human brain: selective expression in gray matter astrocytes, co-localization with connexin43 at gap junctions and late developmental appearance. Neuroscience 1999; 88: 447–68.

65. Ezan P, André P, Cisternino S, Saubaméa B, Boulay A-C, Doutremer S et al. Deletion of Astroglial Connexins Weakens the Blood–Brain Barrier. J Cereb Blood Flow Metab 2012; 32: 1457–1467.

66. De Bock M, Vandenbroucke RE, Decrock E, Culot M, Cecchelli R, Leybaert L. A new angle on blood–CNS interfaces: A role for connexins? FEBS Lett 2014; 588: 1259–1270.

67. Rouach N, Koulakoff A, Abudara V, Willecke K, Giaume C. Astroglial Metabolic Networks Sustain Hippocampal Synaptic Transmission. Science (80-) 2008; 322: 1551–1555.

68. O’Leary L, Belliveau C, Davoli M-A, Mechawar N. Program No. 042.01. Regional characterization of vimentin-immunoreactive astrocytes in the human brain. Neurosci Meet Planner San Diego, CA Soc Neurosci 2018 Online 2018.

69. Jeanson T, Pondaven A, Ezan P, Mouthon F, Charvériat M, Giaume C. Antidepressants Impact Connexin 43 Channel Functions in Astrocytes. Front Cell Neurosci 2015; 9: 495.

70. Kuhlmann T, Remington L, Maruschak B, Owens T, Brück W. Nogo-A is a reliable oligodendroglial marker in adult human and mouse CNS and in demyelinated lesions. J Neuropathol Exp Neurol 2007; 66: 238–246.

71. Laird DW. The gap junction proteome and its relationship to disease. Trends Cell Biol 2010; 20: 92–101.

72. Dianati E, Poiraud J, Weber-Ouellette A, Plante I. Connexins, E-cadherin, Claudin-7 and β-catenin transiently form junctional nexuses during the post-natal mammary gland development. Dev Biol 2016; 416: 52–68.

73. Dbouk HA, Mroue RM, El-Sabban ME, Talhouk RS. Connexins: a myriad of functions extending beyond assembly of gap junction channels. Cell Commun Signal 2009; 7: 4.

74. Martins-Marques T, Anjo SI, Pereira P, Manadas B, Girão H. Interacting Network of the Gap Junction (GJ) Protein Connexin43 (Cx43) is Modulated by Ischemia and Reperfusion in the Heart. Mol Cell Proteomics 2015; 14: 3040–55.

75. Talhouk RS, Mroue R, Mokalled M, Abi-Mosleh L, Nehme R, Ismail A et al. Heterocellular interaction enhances recruitment of α and β-catenins and ZO-2 into functional gap-junction complexes and induces gap junction-dependant differentiation of mammary epithelial cells. Exp Cell Res 2008; 314: 3275–3291.

76. Fujimoto K, Nagafuchi A, Tsukita S, Kuraoka A, Ohokuma A, Shibata Y. Dynamics of connexins, E-cadherin and alpha-catenin on cell membranes during gap junction formation. J Cell Sci 1997; 110 (Pt 3): 311–22.

77. Hunter AW, Barker RJ, Zhu C, Gourdie RG. Zonula occludens-1 alters connexin43 gap junction size and organization by influencing channel accretion. Mol Biol Cell 2005; 16: 5686–98.

78. Singh D, Solan JL, Taffet SM, Javier R, Lampe PD. Connexin 43 Interacts with Zona Occludens-1 and −2 Proteins in a Cell Cycle Stage-specific Manner. J Biol Chem 2005; 280: 30416–30421.

79. Kojima T, Sawada N, Chiba H, Kokai Y, Yamamoto M, Urban M et al. Induction of Tight Junctions in Human Connexin 32 (hCx32)-Transfected Mouse Hepatocytes: Connexin 32 Interacts with Occludin. Biochem Biophys Res Commun 1999; 266: 222–229.

80. Kojima T, Kokai Y, Chiba H, Yamamoto M, Mochizuki Y, Sawada N. Cx32 but Not Cx26 Is Associated with Tight Junctions in Primary Cultures of Rat Hepatocytes. Exp Cell Res 2001; 263: 193–201.

81. Denninger AR, Breglio A, Maheras KJ, LeDuc G, Cristiglio V, Demé B et al. Claudin-11 Tight Junctions in Myelin Are a Barrier to Diffusion and Lack Strong Adhesive Properties. Biophys J 2015; 109: 1387–97.

82. Gow A, Southwood CM, Li JS, Pariali M, Riordan GP, Brodie SE et al. CNS myelin and sertoli cell tight junction strands are absent in Osp/claudin-11 null mice. Cell 1999; 99: 649–59.

83. Morita K, Sasaki H, Fujimoto K, Furuse M, Tsukita S. Claudin-11/OSP-based tight junctions of myelin sheaths in brain and Sertoli cells in testis. J Cell Biol 1999; 145: 579–88.

84. Schubert A-L, Schubert W, Spray DC, Lisanti MP. Connexin family members target to lipid raft domains and interact with caveolin-1. Biochemistry 2002; 41: 5754–64.

85. Langlois S, Cowan KN, Shao Q, Cowan BJ, Laird DW. Caveolin-1 and −2 Interact with Connexin43 and Regulate Gap Junctional Intercellular Communication in Keratinocytes. Mol Biol Cell 2008; 19: 912–928.

86. Giepmans B. Gap junctions and connexin-interacting proteins. Cardiovasc Res 2004; 62: 233–245.

87. Butkevich E, Hülsmann S, Wenzel D, Shirao T, Duden R, Majoul I. Drebrin Is a Novel Connexin-43 Binding Partner that Links Gap Junctions to the Submembrane Cytoskeleton. Curr Biol 2004; 14: 650–658.

88. Ursitti JA, Petrich BG, Lee PC, Resneck WG, Ye X, Yang J et al. Role of an alternatively spliced form of alphaII-spectrin in localization of connexin 43 in cardiomyocytes and regulation by stress-activated protein kinase. J Mol Cell Cardiol 2007; 42: 572–81.

89. Toyofuku T, Yabuki M, Otsu K, Kuzuya T, Hori M, Tada M. Direct association of the gap junction protein connexin-43 with ZO-1 in cardiac myocytes. J Biol Chem 1998; 273: 12725–31.

90. Majoul I, Shirao T, Sekino Y, Duden R. Many faces of drebrin: from building dendritic spines and stabilizing gap junctions to shaping neurite-like cell processes. Histochem Cell Biol 2007; 127: 355–361.

91. Ernst C, Nagy C, Kim S, Yang JP, Deng X, Hellstrom IC et al. Dysfunction of astrocyte connexins 30 and 43 in dorsal lateral prefrontal cortex of suicide completers. Biol Psychiatry 2011; 70: 312–319.

92. Bronstein JM, Tiwari-Woodruff S. Tight Junctions in CNS Myelin. In: Tight Junctions. Springer US: Boston, MA, 2006, pp 196–205.

93. Brites D, Fernandes A. Neuroinflammation and Depression: Microglia Activation, Extracellular Microvesicles and microRNA Dysregulation. Front Cell Neurosci 2015; 9: 476.

94. Torres-Platas SG, Cruceanu C, Chen GG, Turecki G, Mechawar N. Evidence for increased microglial priming and macrophage recruitment in the dorsal anterior cingulate white matter of depressed suicides. Brain Behav Immun 2014; 42: 50–9.

95. Devorak J, Torres-Platas SG, Davoli MA, Prud’homme J, Turecki G, Mechawar N. Cellular and Molecular Inflammatory Profile of the Choroid Plexus in Depression and Suicide. Front Psychiatry 2015; 6. doi:10.3389/fpsyt.2015.00138.

96. Rochfort KD, Cummins PM. Cytokine-mediated dysregulation of zonula occludens-1 properties in human brain microvascular endothelium. Microvasc Res 2015; 100: 48–53.

97. Van Itallie CM, Fanning AS, Holmes J, Anderson JM. Occludin is required for cytokine-induced regulation of tight junction barriers. J Cell Sci 2010; 123: 2844–52.

98. Capaldo CT, Farkas AE, Hilgarth RS, Krug SM, Wolf MF, Benedik JK et al. Proinflammatory cytokine-induced tight junction remodeling through dynamic self-assembly of claudins. Mol Biol Cell 2014; 25: 2710–9.

99. Menard C, Pfau ML, Hodes GE, Kana V, Wang VX, Bouchard S et al. Social stress induces neurovascular pathology promoting depression. Nat Neurosci 2017; 20: 1752–1760.

100. Castellano P, Eugenin EA. Regulation of gap junction channels by infectious agents and inflammation in the CNS. Front Cell Neurosci 2014; 8: 122.

101. Olympiou M, Sargiannidou I, Markoullis K, Karaiskos C, Kagiava A, Kyriakoudi S et al. Systemic inflammation disrupts oligodendrocyte gap junctions and induces ER stress in a model of CNS manifestations of X-linked Charcot-Marie-Tooth disease. Acta Neuropathol Commun 2016; 4: 95.

102. Papaneophytou CP, Georgiou E, Karaiskos C, Sargiannidou I, Markoullis K, Freidin MM et al. Regulatory role of oligodendrocyte gap junctions in inflammatory demyelination. Glia 2018; 66: 2589–2603.

103. Markoullis K, Sargiannidou I, Schiza N, Hadjisavvas A, Roncaroli F, Reynolds R et al. Gap junction pathology in multiple sclerosis lesions and normal-appearing white matter. Acta Neuropathol 2012; 123: 873–886.

104. Markoullis K, Sargiannidou I, Schiza N, Roncaroli F, Reynolds R, Kleopa KA. Oligodendrocyte Gap Junction Loss and Disconnection From Reactive Astrocytes in Multiple Sclerosis Gray Matter. J Neuropathol Exp Neurol 2014; 73: 865–879.

105. Drevets WC, Savitz J, Trimble M. The subgenual anterior cingulate cortex in mood disorders. CNS Spectr 2008; 13: 663–81.

106. Teicher MH, Samson J a., Anderson CM, Ohashi K. The effects of childhood maltreatment on brain structure, function and connectivity. Nat Rev Neurosci 2016; 17: 652–666.

107. Sarrouilhe D, Mesnil M, Dejean C. Targeting Gap Junctions: New Insights in the Treatment of Major Depressive Disorder. Curr Med Chem 2018; 25. doi:10.2174/0929867325666180327103530.

